# Developmental Circadian Disruption Alters Placental Signaling in Mice

**DOI:** 10.1101/2021.04.21.440521

**Authors:** Danielle A. Clarkson-Townsend, Katie L. Bales, Karen E. Hermetz, Amber A. Burt, Machelle T. Pardue, Carmen J. Marsit

**Affiliations:** Gangarosa Department of Environmental Health, Rollins School of Public Health, Emory University, Atlanta, GA, USA; Center for Visual and Neurocognitive Rehabilitation, Atlanta VA Healthcare System, Decatur, GA, USA; Department of Ophthalmology, Emory University, Atlanta, GA, USA; Department of Biomedical Engineering, Georgia Institute of Technology and Emory University, Atlanta, GA, USA

**Keywords:** placenta, developmental chronodisruption, circadian disruption, developmental programming, DOHaD

## Abstract

Circadian disruption has been largely overlooked as a developmental exposure. The placenta, a conduit between the maternal and fetal environments, may relay circadian cues to the fetus. We have previously shown that developmental chronodisruption causes visual impairment and increased retinal microglial and macrophage marker expression. Here, we investigated the impacts of environmental circadian disruption on fetal and placental outcomes in a C57BL/6J mouse (*Mus musculus*) model. Developmental chronodisruption had no effect on embryo count, placental weight, or fetal sex ratio. When measured with RNAseq, mice exposed to developmental circadian disruption (CD) had differential placental expression of several transcripts including *Serpinf1*, which encodes pigment-epithelium derived factor (PEDF). Immunofluorescence of microglia/macrophage markers, Iba1 and CD11b, also revealed significant upregulation of immune cell markers in CD-exposed placenta. Our results suggest that *in utero* circadian disruption enhances placental immune cell expression, potentially programming a pro-inflammatory tissue environment that increases the risk of chronic disease in adulthood.

## INTRODUCTION

Environmental light exposure has changed rapidly over the last century with the introduction of electric lighting. One of the consequences of the modern light environment is circadian disruption, or misalignment between the internal temporal system and external cues(1). Circadian disruption can promote the development of chronic diseases, such as diabetes and dyslipidemia(2-5); night shift work is even categorized by the International Agency for Research on Cancer as a Group 2A carcinogen, “probably carcinogenic to humans”(6). However, little is known how circadian disruption affects fetal development.

The Developmental Origins of Health and Disease (DOHaD) hypothesis grew out of research on *in utero* undernutrition and later life risk of cardiometabolic disease(7, 8). These studies found that infants born with low birthweight or small for their gestational age (SGA) had an increased risk of heart disease and stroke as adults(9-13). Later, the Dutch Hunger Winter cohort revealed epigenetic(14) and transgenerational(15) effects of *in utero* exposure to famine on offspring. DOHaD research has grown to encompass exposure to early life stress and pollutants(16), such as endocrine disrupting compounds, and outcomes related to neurological and hormonal programming. Light can also act as an endocrine disruptor(17); however, the influence of light exposure on developmental programming has not yet been widely assessed in DOHaD studies.

We have previously shown that developmental chronodisruption in mice (via environmental light) from embryonic day 0 until weaning at 3 weeks of age has lasting effects on visual and metabolic outcomes of adult offspring; in particular, mice exposed to developmental circadian disruption have increased expression of retinal microglia and macrophage markers accompanied with impaired visual function(18). The placenta, a neuroendocrine organ, regulates *in utero* growth, including fetal neuronal growth. Communication between the placenta and fetal brain, termed the placenta-brain axis(19), influences neurodevelopment. The immune system plays an important role in the placenta-brain axis, and activation of placental immune signals can influence development of fetal immune cells, such as microglia, in the fetal brain(20, 21). Therefore, we investigated the impacts of developmental circadian disruption on overall gene expression and immune cell phenotypes in the placenta. To do this, we exposed pregnant mice to developmental chronodisruption and measured fetal and placenta outcomes (count, weight, sex ratio), placental gene expression (RNAseq), and placental expression of immune cell markers CD11b and Iba1 (immunofluorescence).

## MATERIALS and METHODS

### Ethical approval

All experimental procedures were approved (#V008-19) by the Institutional Animal Care and Use Committee of the Atlanta Veterans Affairs Healthcare System in facilities that are accredited by the Association for the Assessment and Accreditation of Laboratory Animal Care International (AAALAC).

### Animal handling and experimental design

Wildtype female (∼3-4 weeks old) C57BL/6J mice (*Mus musculus*) were ordered from Jackson Laboratories (Bar Harbor, ME, USA); wildtype male C57BL/6J mice were ordered or bred in-house from mice from Jackson Laboratories. Males for breeding were singly housed whereas female breeders were co-housed in large (6”x9”x18”) wire-top shoebox cages in standard conditions (*ad libitum* chow (Teklad Rodent Diet 2018 irradiated 2918, Envigo Teklad, Madison, WI, USA), 12:12 lighting) and checked daily for well-being. After a 2 week acclimation period, naïve females were randomized to either control light (CL, 12:12 light:dark) or a chronodisruption (CD) light paradigm, consisting of weekly inversions of the photoperiod(18, 22, 23). Light intensity was standardized across groups to be ∼50-400 lux (Dual-range light meter 3151CC, Traceable, Webster, TX, USA), with darkest areas at the bottom of the cage under the food holder and brightest areas near the top of the cage. Females were exposed to light treatments for 4 weeks prior to timed breeding; during aligned light schedules, representative females from each light treatment group were introduced to the male’s cage in the afternoon; females were checked for plugs and returned to their home cages after 2 days.

Females were weighed several days later to confirm pregnancy; if not pregnant, they were placed with the same male the following week for further rounds of pairing for up to 4 more weeks of pre-pregnancy light treatment. Dams remained in CD or CL light treatments until tissue collection at gestational day 15.5 (E15.5). While placental tissue collection was timed to be the estimated E15.5 and mouse pairings occurred in a restricted time window, we did not evaluate vaginal cytology or use *in vitro* fertilization, and it is therefore possible that embryonic age varied by a day.

### Tissue collection

Pregnant mice (E15.5) were sacrificed with compressed CO_2_ gas anesthesia, followed by cervical dislocation and rapid decapitation for truncal blood collection between 9AM-11AM (ZT3-5); within this range, tissue collection time did not substantially differ between CL and CD groups. Position of each placental sample within the uterine horns, placental wet weight, and reabsorptions were recorded and placentae immediately dissected out after removing uterine tissue. Placental tissue samples were snap-frozen in liquid nitrogen and stored at -80°C until further processing for RNA isolation or preserved in 10% neutral buffered formalin for histological and immunohistochemical analyses. Fetal tail samples were also collected, snap frozen, and stored at -80°C until later use for sex determination. Samples from 3 dams, all from the CD group, were excluded due to noted quality issues during collection; for example, in 2 mice, all of the embryos in a uterine horn exhibited blood clots and discoloration. Samples from a total of 12 dams, 6 CL and 6 CD, were included in the analysis.

### RNA isolation, sequencing, alignment, and generation of count data

Prior to placental RNA isolation, all fetal tail tissue samples were lysed and RNA extracted using the Qiagen Allprep DNA/RNA Mini Kit according to manufacturer’s instructions and *Sry* gene expression measured via PCR to determine sex (SryFWD: 5’ – TGG GAC TGG TGA CAA TTG TG -3’ and SryREV : 5’ – GAG TAC AGG TGT GCA GCT CT-3’). Samples with faint bands were re-run. For RNA sequencing, placental samples without any noted collection quality issues were randomly selected and matched on sex when possible (quality samples of both sexes were not available for each dam). Two samples from each dam were chosen, for a total of 24 placenta samples, 12 from each light treatment group, and DNA and RNA isolated using the Qiagen Allprep DNA/RNA Mini Kit according to manufacturer’s instructions. RNA quality was measured using the Agilent 2100 Bioanalyzer with Agilent RNA 6000 Nano kit (cat# 5067-1511) following manufacturer’s instructions and RNA concentrations measured with a Thermo Scientific NanoDrop spectrophotometer. All samples had RIN scores ≥ 9. Placental RNA samples (n=24) were sent to the Emory Genomics core for PolyA RNA sequencing performed at 30M read depth. FastQC was performed to check read quality and fastq files aligned to the C57 mouse genome (Ensemble assembly GRCm38.p6) with STAR v2.7 using default settings. Read counts were derived using the “quantmode” command in STAR. Raw sequencing data FastQ files, processed gene count data, and sample information have been deposited in GEO (accession number GSE169266). Code for sample alignment and processing, as well as gene count data, are available at: https://github.com/dclarktown/CD_mice_placenta (DOI: 10.5281/zenodo.4536522).

### Differential expression (DE) analysis

Count data were read into R (version 3.2) and analyzed for differential expression (DE) using DESeq2(24). The original 53,801 transcripts measured were limited to transcripts that had at least 1 count in 10% of samples, leaving a total of 14,739 transcripts for analysis. To confirm sex of samples, samples were also evaluated for high expression of *Xist* mRNA, indicative of female sex. Of the 24 samples, 1 sample was mismatched for sex (sample #9, 1009) and edited to the correct sex. After 1 sample was found to be an outlier driving many of the DE results (sample #12, 1012), it was dropped from the analysis. The DE analysis of the remaining 23 samples adjusted for sex and the first surrogate variable, with developmental light treatment group as the main exposure. The first surrogate variable was computed using the *sva* package(25) and “be” method with 200 iterations. Results were adjusted for false discovery rate using the Benjamini and Hochberg (BH) method and considered significant if q<0.05.

### Pathway analysis

Transcript enrichment for differentially expressed genes was performed using EnrichR(26) among the Mouse Gene Atlas, ChEA 2016, KEGG 2019 Mouse, and GO 2018 (Biological Process, Molecular Function, Cellular Component) databases. Results were adjusted for multiple comparisons using the Benjamini-Hochberg (BH) method and considered significant if q<0.05.

### Placental immunofluorescence measurement and quantification

Fresh placenta samples were fixed in 10% neutral buffered formalin overnight at 4°C and then cryoprotected the following day in 30% sucrose after washing with 1x PBS. Samples were embedded and frozen in optimal cutting temperature compound and sliced into 7-µm-thick sections. Placental sections were blocked (with 0.1% Triton X-100) and incubated with primary antibodies in 5% normal donkey serum in PBS before washing with PBS. Primary antibody incubations using Iba1 (ab178847; 1:100; Abcam) and CD11b (14-0112-82; 1:100; Invitrogen) were performed for 16-24 hours at 4°C and secondary antibody incubations were performed for 1 hour at room temperature using Alexa Fluor 488 Donkey anti-mouse IgG (A-21202; 1:500) and Alexa Fluor 647-conjugated Donkey anti-rabbit IgG (A-31573; 1:500). Tissue nuclei were visualized with nuclear stain 4′,6-diamidino-2-phenylindole (DAPI, 62247; Thermo Fisher Scientific). Coverslips were mounted using Prolong Gold (P36934; Thermo Fisher Scientific). Placental tissue (n=4-6 mice/group; 3 images per sample, averaged for the analysis) was imaged with an Olympus Fluoview1000 confocal microscope (Center Valley, PA) using a 20x objective and a Lumenera INFINITY 1-3C USB 2.0 Color Microscope camera (Spectra Services, Ontario, NY). All images were processed and quantified using ImageJ software by a researcher masked to treatment group.

### Statistical analysis and data availability

Unless otherwise noted, weight, embryo number, placental weight, sex ratio, and immunofluorescence data were all analyzed with Student’s 2-tailed unpaired t-tests and considered significant if p<0.05. Statistical tests were performed in Prism version 9.0.0. Statistics for placental gene expression analyses are described in the previous sections. All data and code used for the analyses are available at: https://github.com/dclarktown/CD_mice_placenta (DOI: 10.5281/zenodo.4536522), except for the raw sequencing data which has been deposited in GEO (accession number GSE169266).

## RESULTS and DISCUSSION

Here, we investigated whether circadian disruption led to gene expression and immunologic changes in the placenta. We have previously shown that developmental CD light treatment alters programming of the visual system in offspring(18). CD females did not differ in pre-pregnancy weight (Student’s unpaired 2-tailed t-test, t=0.83, df=10, p=0.43, **Figure 1A**) or pregnancy weight at tissue collection (Student’s unpaired 2-tailed t-test, t=0.72, df=10, p=0.49, **Figure 1B**) compared to CL females. There were also no differences in embryo count, fetal sex ratio, and placental weight (**Figure 1C-F**), consistent with previous findings(27) in a rat model that additionally found no change in fetal weight or placental:fetal weight ratio. Genetic models of developmental circadian disruption have found similar null results; knockout of *Bmal1* (*Arntl*), a core circadian clock gene, in fetal tissue does not alter embryo number or fetal or placental weight(28), whereas knockout in parental male or female tissue causes infertility(29). However, the exclusion of 2 CD dams from the analysis due to discoloration and blood clots throughout one uterine horn may have biased results towards a more conservative measure of effect.

**Figure 1.**
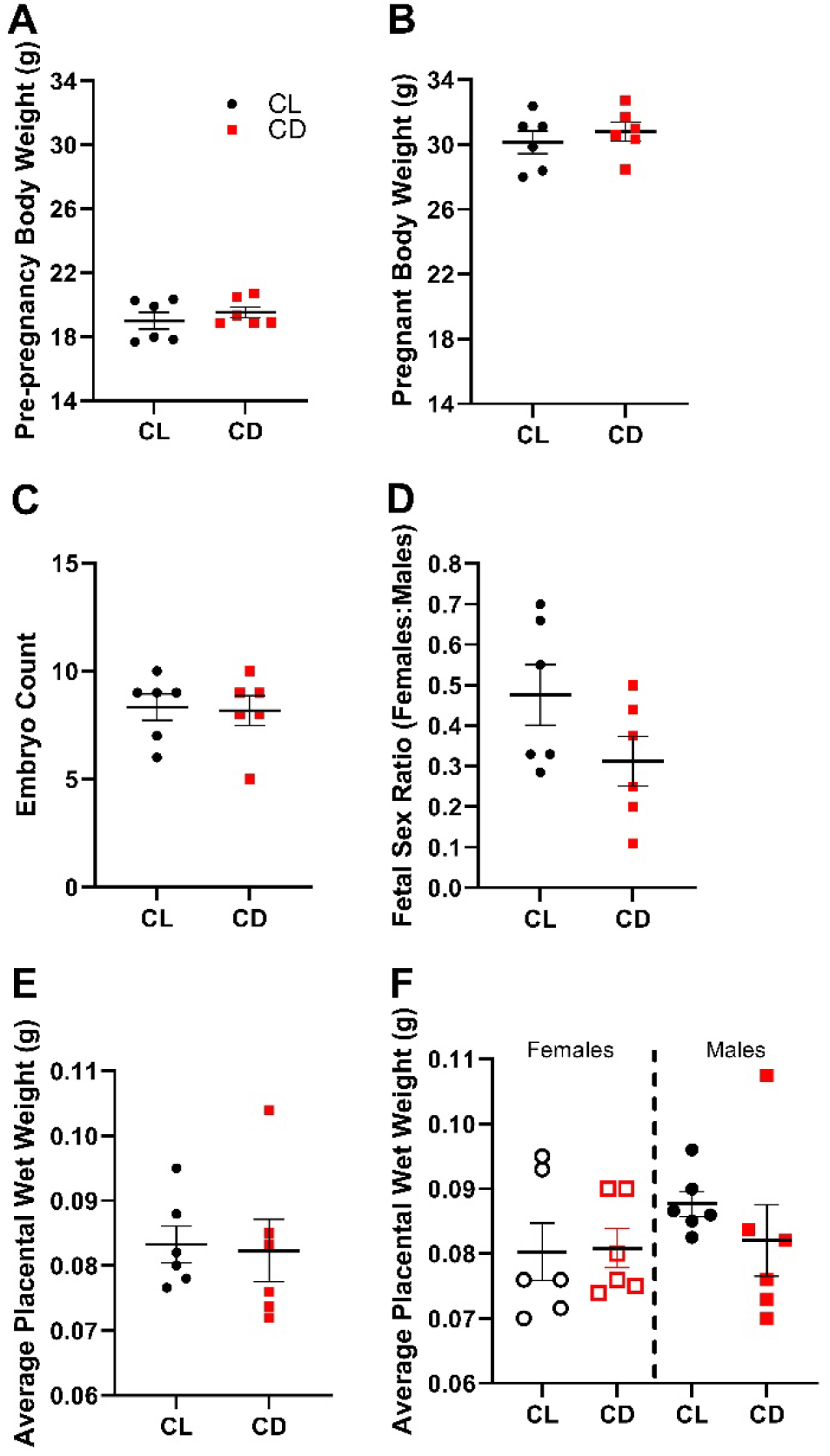
Light treatment did not alter dam or fetal outcomes. Body weight (grams) of female mice (CL n=6, CD n=6) from which placental samples were collected (**A**) just prior to pairing for timed breeding (Student’s unpaired 2-tailed t-test, t=0.83, df=10, p=0.43) and (**B**) when pregnant at E15.5 just prior to tissue collection (Student’s unpaired 2-tailed t-test, t=0.72, df=10, p=0.49). (**C**) Number of viable embryos per dam counted within the uterine horns (Student’s 2-tailed unpaired t-test, t=0.18, df=10, p=0.86). (**D**) Sex ratio of viable embryos per dam, as determined by PCR of fetal tail snip and subset confirmed by RNA sequencing (Student’s 2-tailed unpaired t-test, t=1.67, df=10, p=0.12). (**E**) Average placental wet weight (grams) per dame (Student’s 2-tailed unpaired t-test, t=0.16, df=10, p=0.87). (**F**) Average placental wet weight (grams) per dam, stratified by sex (1-way ANOVA, F (3, 20) = 0.73, p=0.55). All data are presented as mean ± SEM.

Placentas were collected at the late stage of pregnancy and sequenced for gene expression. Among the most highly expressed transcripts across all placenta samples (regardless of exposure) were *Tpbpa, Prl3b1, Tpbpb, Psg21, Prl8a9*, and *Psg23*, gene expression typical of trophoblasts(30). The EnrichR pathway analysis of the top 100 most highly expressed placental genes indicated enrichment for mouse placental tissue (q<0.05, **Supplemental File 1**) in the Mouse Gene Atlas database, as expected, and, interestingly, for the CLOCK, NELFA, and HSF1 transcription factors in the ChEA database. CLOCK is a core component of the circadian clock, and as mediator of the maternal and fetal environments, the placenta may function as a peripheral oscillator; we have previously shown that placental gene expression varies seasonally(31), which suggests sensitivity to seasonal environmental exposures such as light and temperature. Top KEGG pathways were: “antigen processing and presentation”, “protein processing in endoplasmic reticulum”, “lysosome” and “HIF-1 signaling pathway”; likewise, top GO pathways were related to immune signaling and protein processing, with terms such as “ATF6-mediated unfolded protein response”, “neutrophil degranulation”, “collagen binding”, “secretory granule lumen”, and “focal adhesion” (**Supplemental File 1**).

Principle component analysis of the placental samples revealed relative overlap between the CL and CD groups (**Figure 2A**). This pattern was not explained by sample position within uterine horn, sample collection time, sex ratio, or RNA quality, and samples from the same dam did not necessarily cluster together. The DE analysis between male and female placental tissue (adjusting for light treatment) resulted in 77 sex-specific placental transcripts (q<0.05, **Figure 2B, Supplemental File 2**). A number of these genes were strikingly different; *Xist*, a non-coding RNA that silences the extra X-chromosome in females and can be used to identify fetal sex(32), was highly expressed in female placenta. *Ddx3y, Eif2s3y, Kdm5d*, and *Uty* were all highly expressed in male placenta and have previously been reported as male-specific placental genes(33, 34); these genes could arguably also be used to identify fetal sex. Sex-specific placental gene expression (n=113 q<0.1) also displayed enrichment for pathways related to lipid, retinoid, and cholesterol metabolism in the KEGG and GO term databases, suggesting sex-specific regulation of these processes in the placenta (**Supplemental File 3**). Interestingly, studies of maternal malnutrition and high fat diet exposure have uncovered sex-specific placental(35-37) and phenotypic outcomes in the offspring(38, 39). These results support investigation of these pathways in sex-specific development in future studies.

**Figure 2.**
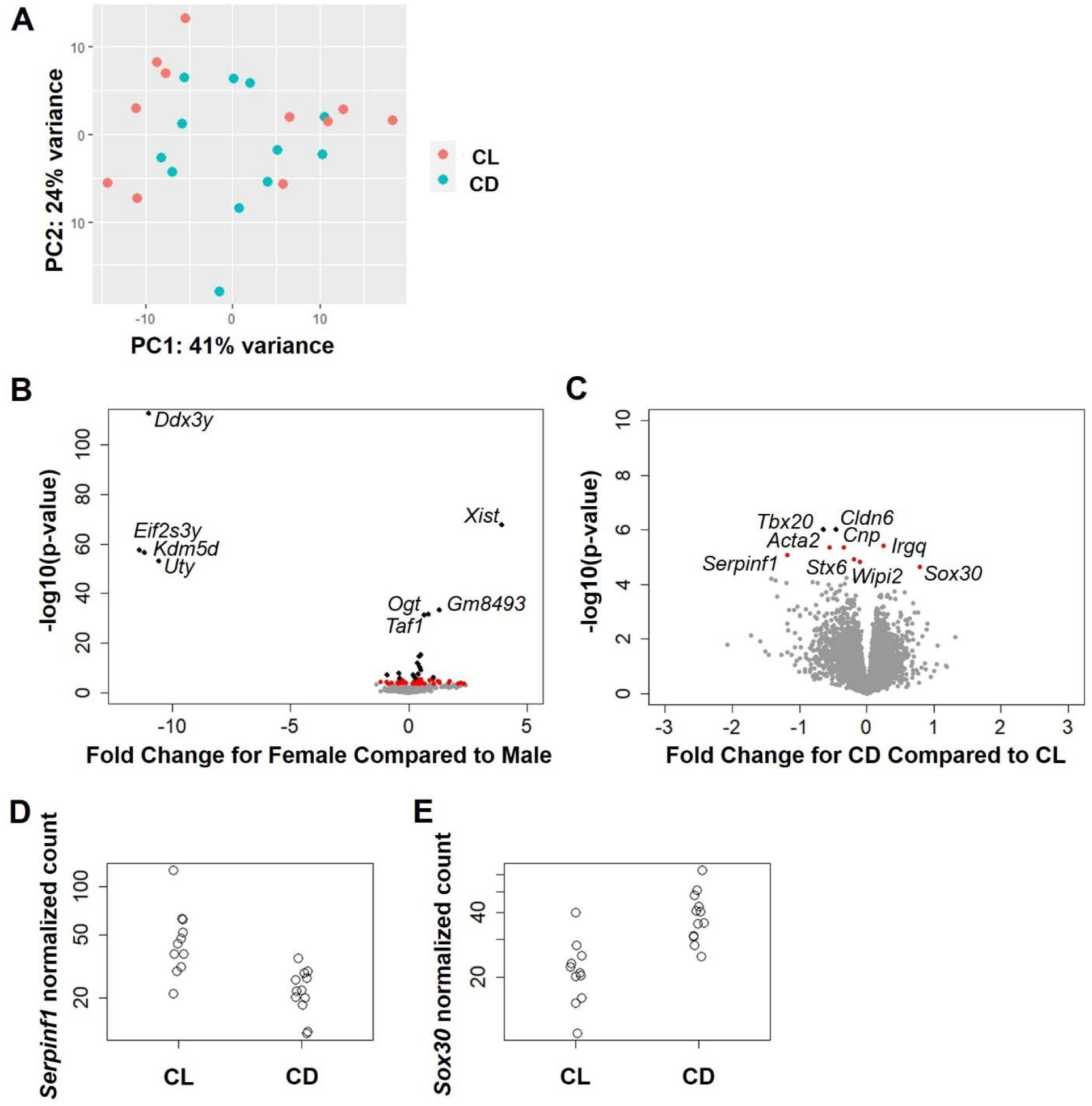
Placental gene expression varies by sex and light treatment group. (**A**) PCA plot of first 2 principal components comparing treatment groups shows general overlap between CL and CD groups. (**B**) Volcano plot of differential placental gene expression by sex (adjusting for light treatment group and first surrogate variable). Male is the reference group, so transcripts with decreased expression in females (or, conversely, increased expression in males) are located to the left of 0, while transcripts with increased expression in females (or, conversely, decreased expression in males) are located to the right of the 0. Black dots denote Bonferroni-significant transcripts (n=22 p<0.05), red dots denote BH-significant transcripts (n=77 q<0.05), and grey dots denote non-significant transcripts. The top differentially expressed genes are plotted with their respective gene names. (**C**) Volcano plot of differential placental gene expression by treatment (adjusting for sex and first surrogate variable). CL is the reference group, so transcripts with decreased expression in CD (or, conversely, increased expression in CL) are located to the left of the 0, while transcripts with increased expression in CD (or, conversely, decreased expression in CL) are located to the right of the 0. Black dots denote Bonferroni-significant transcripts (n=2 p<0.05), red dots denote BH-significant transcripts (n=9 q<0.05) and grey dots denote non-significant transcripts. Plots of raw normalized count data for (**D**) *Serpinf1* and (**E**) *Sox30* by treatment group.

Few transcripts exhibited large differences between light treatment groups (**Figure 2C**). However, of the differentially expressed genes (n=9 q<0.05, **Supplemental File 4**), *Serpinf1, Tbx20, Acta2, Cldn6, Cnp, Stx6*, and *Wipi2* had decreased expression while *Sox30* and *Irgq* had increased expression in CD placenta (**Figure 2C-E**). Pathway analysis revealed that differentially expressed genes were similar to gene expression in osteoblasts in the Mouse Gene Atlas database (**Supplemental File 5**). While there was no enrichment for specific transcription factors within the ChEA database, differentially expressed genes were enriched for “cholesterol metabolism” in the KEGG database and terms related to tissue development, adhesion, and cytoplasmic projection in the GO databases. It is perhaps surprising that we did not uncover large differences in placental gene expression between light treatment groups. However, it is possible the small sample size limited the ability to measure more subtle differences in gene expression, especially if such differences occurred in placental cell subpopulations, such as immune cells.

Placenta from CD-exposed dams revealed significantly increased expression of Iba1 and CD11b microglial/macrophage markers than placenta from dams housed in CL conditions (p=0.027 and p=0.038, respectively, **Figure 3**). These results align with the finding of decreased *Serpinf1* (which encodes pigment epithelium-derived factor (PEDF)) expression in CD placenta. A neurotrophic factor with many roles(40), PEDF inhibits macrophage inflammatory processes(41), which may have contributed to the increased CD11b and Iba1 marker expression in CD placenta. These data also coincide with our findings of an increased retinal inflammatory response and reduced visual function within mice developmentally exposed to CD(18). The immune system and inflammation govern many of the health outcomes caused by chronic circadian disruption(42-44). For example, night shift workers were found to have greater amounts of immune cells, such as T cells and monocytes, than non-shift workers(45); likewise, we previously reported hypomethylation in immune-related genes, such as *CLEC16A, SMPD1*, and *TAPBP*, in the placentas of mothers who worked the night shift(46). In rodent studies, chronic circadian disruption increased macrophages and “pro-tumor” CD11b+ MHCII cells(47), altered inflammatory response in the brain(48), and primed the innate immune response to be more pro-inflammatory(49).

**Figure 3.**
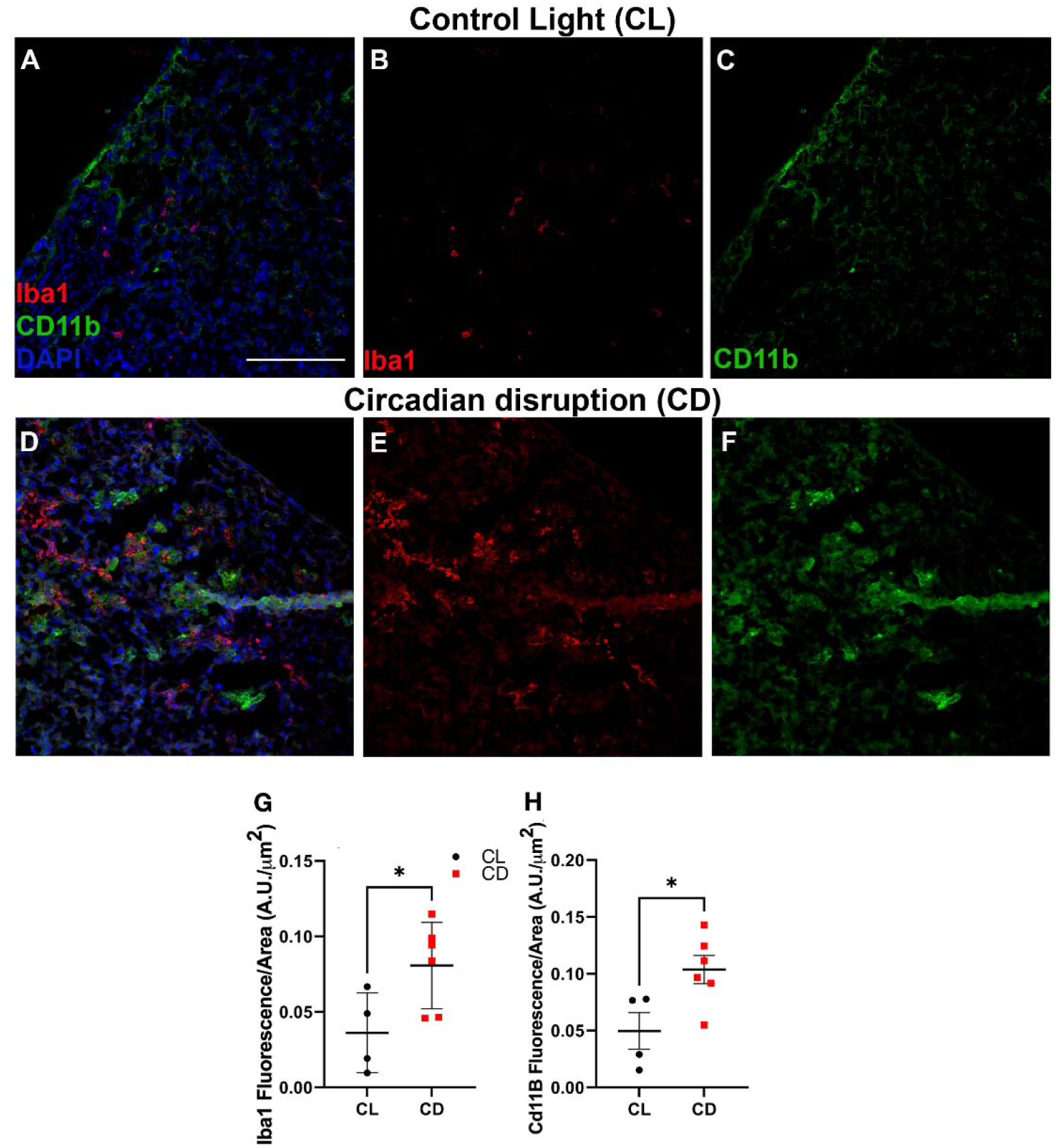
Chronodisruption causes increased macrophage and microglial signaling in the placenta. Placentas from (**A-C**) control light (CL) and mice exposed to (**D-F**) developmental circadian disruption (CD) were labeled for inflammatory markers labeling microglia and macrophages. In placenta from CD mice increased placental (**G**) Iba1 fluorescence (Student’s 2-tailed unpaired t-test, t=2.49, df=8, p=0.038) and increased placental **(H**) CD11b fluorescence (Student’s 2-tailed unpaired t-test, t=2.70, df=8, p=0.027) were detected. CL=4 placenta from different dams, 3 images each; CD=6 placenta from different dams, 3 images each. All data are presented as mean ± SEM, scale bar = 20 microns, and *=p<0.05.

The placenta is the only organ formed by the interaction of both fetal/embryonic and maternal tissues and acts as the interface between both circulatory systems(50). Previous research has found a strong correlation between placental CD11b expression and fetal brain microglial activation(21). In mice and humans, brain and placental macrophages and microglia originally derive from the same source: the fetal yolk sac(51, 52). These progenitor macrophage and microglial cells migrate from the yolk sac to embryonic tissues, where they set up residence; once settled, they are long-lived and able to replenish themselves(53, 54). Our results suggest that developmental light environment affects programming of the placental and fetal immune systems, laying the groundwork for a pro-inflammatory setting later in life. These findings provide novel evidence linking CD with increased placental inflammatory response and highlight the need to evaluate the influence of the light environment on health and disease outcomes in DOHaD studies.

## Acknowledgements

This work was supported by funding from the National Institutes of Health (NIH-NICHD F31 HD097918 [to DACT], NIH-NIEHS T32 ES012870 [to DACT], NIH-NIEHS P30ES019776 [to CJM], NIH-NEI Core Grant P30EY006360) and the Department of Veterans Affairs (Rehabilitation Research and Development Senior Research Career Scientist Award RX003134 [to MTP]. This study was supported in part by the Emory Integrated Genomics Core (EIGC), which is subsidized by the Emory University School of Medicine and is one of the Emory Integrated Core Facilities. Additional support was provided by the Georgia Clinical & Translational Science Alliance of the National Institutes of Health under Award Number UL1TR002378. The sequencing for this study was supported in part by the Emory Molecules to Mankind (M2M) program, funded by the Burroughs Wellcome Fund. The study sponsors did not have any role in the study design, collection, analysis, interpretation of the data, writing of the report, or the decision to submit the paper for publication.

## Author Contributions

D. Clarkson-Townsend designed the study, performed the experiments, analyzed the data, and wrote and edited the manuscript. K. Bales stained, imaged, quantified, and analyzed placenta samples for immunofluorescence and contributed to writing and editing of the manuscript. K. Hermetz contributed to sample preparation and measurement for mRNA sequencing and editing of the manuscript. A. Burt contributed to genome alignment of sequencing data and editing of the manuscript. M. Pardue provided experimental resources and contributed to experimental design and editing of the manuscript. C. Marsit contributed to experimental design and editing of the manuscript.

## Abbreviations

BH: Benjamini and Hochberg
CL: control light
CD: circadian disruption
DOHaD: Developmental Origins of Health and Disease

## Notes

### Competing Interest Statement

The authors have declared no competing interest.

